# Muscular expression of *pezo-1* differentially contributes to swimming and crawling production in the nematode *C. elegans*

**DOI:** 10.1101/2024.08.13.607367

**Authors:** A Fazyl, E Sawilchik, W Stein, AG Vidal-Gadea

## Abstract

Mechanosensitive PIEZO ion channels are evolutionarily conserved proteins that are widely expressed in neuronal and muscular tissues. This study explores the role of the mechanoreceptor PEZO-1 in the body wall muscles of *Caenorhabditis elegans*, focusing on its influence on two locomotor behaviors, swimming and crawling. Using confocal imaging, we reveal that PEZO-1 localizes to the sarcolemma and plays a crucial role in modulating calcium dynamics that are important for muscle contraction. When we knocked down *pezo-1* expression in striated muscles with RNA interference, calcium levels in head and tail muscles increased. While heightened, the overall trajectory of the calcium signal during the crawl cycle remained the same. While downregulation of *pezo-1* led to an increase in crawling speed, it caused a reduction in swimming speed. Reduction in pezo-1 expression also resulted in the increased activation of the ventral tail muscles, and a disruption of dorsoventral movement asymmetry, a critical feature that enables propulsion in water. These alterations were correlated with impaired swimming posture and path curvature, suggesting that PEZO-1 has different functions during swimming and crawling.

## INTRODUCTION

Locomotion is a fundamental behavior exhibited by virtually all animals, enabling them to interact with their environment, find food, evade predators, and reproduce. Successful locomotion across distinct physical environments, whether crawling on solid surfaces or swimming through liquid, requires overcoming unique challenges posed by each medium. The ability to adapt locomotor patterns to different environments is crucial for survival, and this adaptability is mediated by the neuromuscular system’s capacity to transform neural inputs into appropriate motor outputs.

The relationship between motor input and muscular output is complex and nonlinear. Brezina and colleagues introduced the concept of the Neuromuscular Transform (NMT), demonstrating that neural inputs alone cannot fully explain the behavioral richness exhibited by animals (Brezina et al., 2000a,b; Brezina and Weiss, 2000). They proposed that neuromuscular plasticity plays a significant role in generating adaptive movement, challenging the traditional view of muscles as mere executors of neural commands.

Skeletal muscles, from worms to humans, express a diverse array of mechanoreceptor channels that can alter their muscles’ contractile properties and outputs (Guharay and Sachs, 1984; Hamill and McBride, 1996). Despite their potential important role in muscle function, the specific functions of these mechanoreceptors in muscle physiology and adaptive behaviors are not well understood.

Among the various mechanoreceptors expressed in skeletal muscles, the PIEZO family has emerged as critical mediators of force-sensing across multiple organ systems (Murthy et al., 2017). PIEZO channels have been implicated in processes ranging from touch sensation to vascular tone regulation, yet their specific roles in skeletal muscle function remain underexplored. In the nematode *Caenorhabditis elegans*, which relies on dorsoventral undulations to produce distinct motor gaits for crawling on solid surfaces and swimming through liquid environments (Vidal-Gadea et al., 2011), the PIEZO ortholog *pezo-1* is expressed in a number of tissues including body wall muscles (Park et al., 2024; Millet et al., 2021; Bai et al., 2020; Hughes et al., 2022; Brugman et al., 2022; Noetzel, 2021). However, its role in locomotion has not been fully elucidated (Komandur et al., 2023).

Recent studies have highlighted the importance of proprioceptive feedback in transforming neural inputs into adaptive motor outputs in *C. elegans*. For example, newborn L1 larvae, despite their asymmetrical neural wiring, achieve symmetrical crawling undulations through tonic potentiation of ventral muscles, suggesting that proprioceptive mechanisms play a critical role in locomotor control (Lu et al., 2022). However, most research has focused on neuronal proprioceptors, with limited exploration of the role that mechanoreception within muscles themselves might play in locomotion.

Understanding the role of PEZO-1 in *C. elegans* muscle function is not only essential for deciphering the mechanisms of adaptive locomotion in this organism but also for gaining insights into broader principles of muscle physiology with implications for understanding muscle disorders such as muscular dystrophy in humans.

In this study, we investigate the role of the mechanoreceptor PEZO-1 in the body wall musculature of *C. elegans* during locomotion. By using RNA interference to downregulate *pezo-1* expression specifically in muscle tissues, we examine its differential effects on swimming and crawling behaviors. We propose that the differential expression of mechanoreceptors like *pezo-1* in striated muscles could be a general mechanism by which animals generate diverse and often asymmetrical muscular outputs and behaviors from relatively simple, symmetrical neural inputs. Our findings offer new insights into the interaction between neural and muscular systems in the production of adaptive locomotion and lay the groundwork for further studies on the role of mechanoreceptors in muscle function across species.

## RESULTS

### PEZO-1 is localized to sarcolemma in myocytes

We previously showed that *pezo-1* is expressed in the body wall musculature of *C. elegans* (Hughes et al., 2021). To determine the subcellular localization of PEZO-1 in body wall muscles, we used the AG467 strain expressing an mScarlet-tagged PEZO-1 under its endogenous promoter. Confocal microscopy revealed that PEZO-1 localizes predominantly to the sarcolemma of body wall myocytes, colocalizing with Wheat Germ Agglutinin (WGA) staining of the plasma membrane (Figure 1A, B). This finding is consistent with previous studies suggesting that mechanoreceptors, including PEZO-1, are membrane-bound proteins that transduce mechanical forces into cellular signals (Dong et al., 2010). Our data support the hypothesis that PEZO-1 plays a role in mediating calcium kinetics in these cells, a crucial function for muscle contraction and mechanosensation.

**Figure 1.**
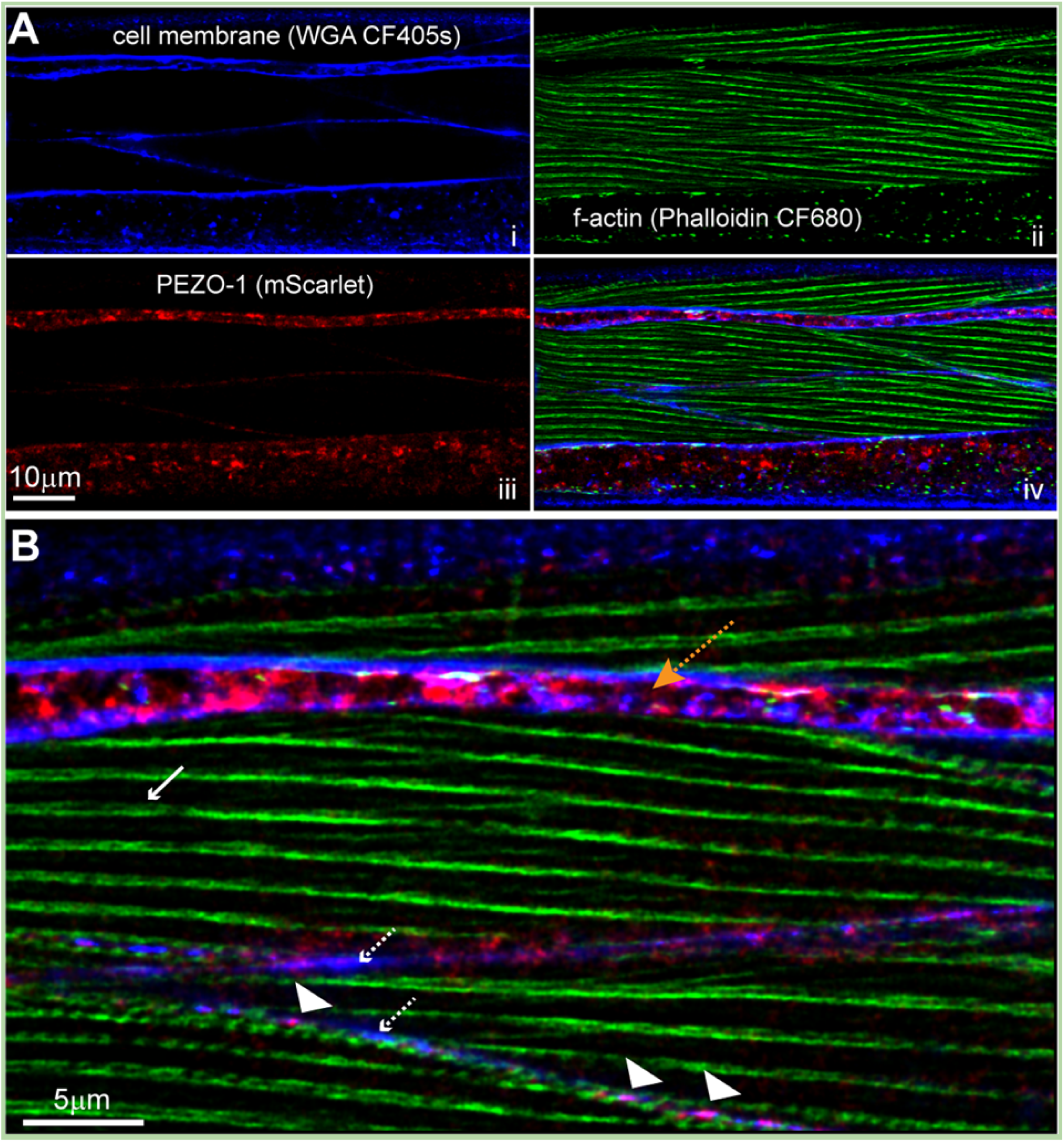
PEZO-1 localization in body wall myocytes. Confocal micrographs of Day-1 AG467 *C. elegans* myocytes expressing the genetically encoded *Ppezo-1::pezo-1::mScarlet* construct. **(A)** Independent channels showing WGA stain of the plasma membrane (blue, i); f-actin filaments in the sarcomere (green, ii); and the genetically encoded mScarlet tagging of the C-terminus of PEZO-1 (red, iii). (iv) Merge of i-iii. **(B)** Magnified view showing PEZO-1 localizing to the plasma membrane. Dashed white arrows indicate the sarcolemma; white solid arrow points to actin filaments; arrowheads indicate PEZO-1; orange arrow points to the hypodermis between muscle quadrants.

### Loss of PEZO-1 causes opposite phenotypes in swimming vs crawling

To assess the functional role of PEZO-1 in locomotion, we employed RNA interference (RNAi) to downregulate *pezo-1* expression in body wall muscles of *C. elegans* (Grishok, 2005). Behavioral analysis using the Tierpsy Tracker software (Javer et al., 2018) revealed that *pezo-1* knockdown produced opposite effects on crawling and swimming speeds. Consistent with our previous findings (Komandur et al., 2023), *pezo-1* RNAi-treated animals displayed a significant increase in crawling speed compared to controls (Figure S1). Interestingly, the same animals exhibited a marked reduction in swimming speed (Figure 2A), a result that was validated using two independent *pezo-1* knockout mutants (Figure 2B). These contrasting effects suggest that PEZO-1 has a context-dependent role in locomotion, potentially related to the differential mechanical demands of crawling on solid surfaces versus swimming in liquid environments.

**Figure 2.**
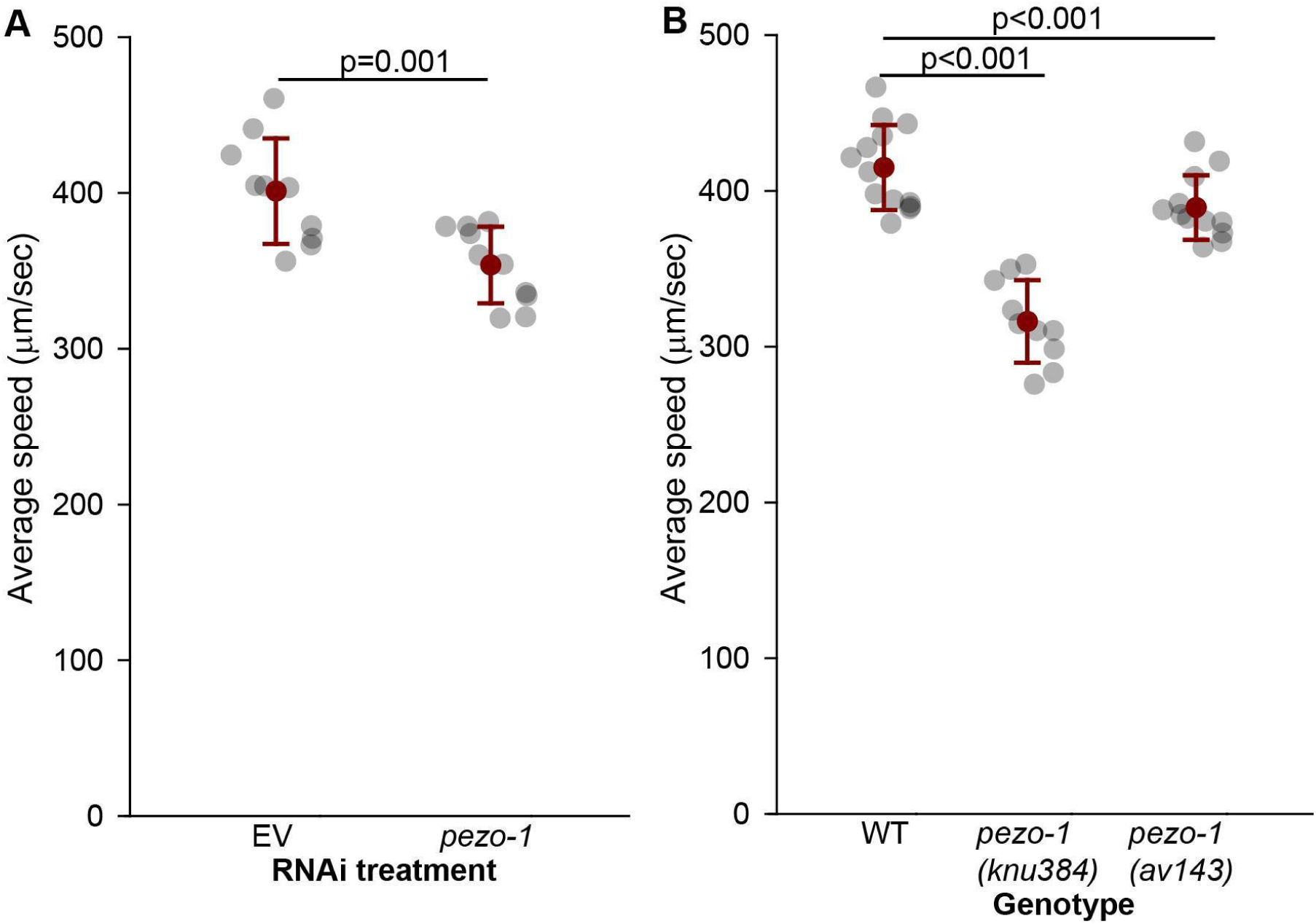
Reduction in *pezo-1* expression decreases swimming speed. **(A)** Average swimming speed of control EV (empty RNAi vector) and *pezo-1* RNAi treated animals was quantified through TIERPSY and resulted in a significant reduction in swimming speed for treated animals. Student t-test. 1-β=0.96. **B.** Validation of findings in A using two *pezo-1* knockout mutants. One way ANOVA (Holm-Sidak) 1-β=1.0. Error bars=SEM. N>10 animals per each (datapoint) assay.

### Crawling is characterized by alternating dorso-ventral body wall muscle activation

We next examined the activation patterns of dorsal and ventral body wall muscles using genetically encoded calcium indicators (GCaMP2) under the *myo-3* promoter. During crawling, control animals exhibited the expected pattern of alternating activation of dorsal and ventral muscles, with peak calcium transients corresponding to peak body bending (Figure 3A), in line with previous reports (Pierce-Shimomura et al., 2008). RNAi-mediated reduction of *pezo-1* expression did not significantly alter this basic pattern but resulted in slightly increased activation of body wall muscles, particularly in the dorsal tail muscles (Figure 3B). Quantification of the area under the curve for muscle activation revealed no significant difference between the overall activation of dorsal versus ventral muscles for both control and *pezo-1* knockdown animals (Figure 3C).

**Figure 3.**
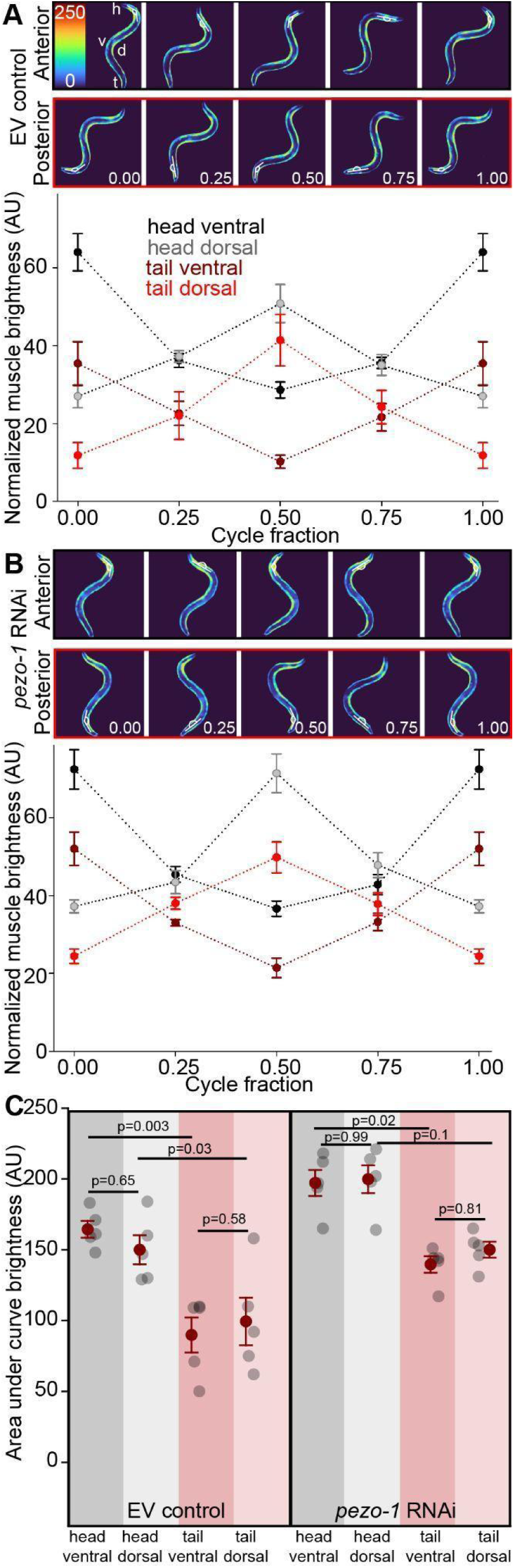
Increased activation of dorsal muscles during crawling following *pezo-1* RNAi. **A.** ZW485 strain expressing GCaMP2 in body wall musculature during crawling showing activation of anterior (top panel) and posterior (bottom panel) contraction cycles. Quantification of muscle brightness during crawling a cycle shows antagonistic activation of dorsal and ventral muscles of the head and tail. **B.** RNAi-mediated reduction in *pezo-1* expression in muscles does not alter the general pattern of muscle activation. **C.** Quantification of area under the curve for A and B shows that reduction in *pezo-1* expression elevates the relative activation of dorsal tail muscles. For L4440 controls (left): One way ANOVA (Holm-Sidak) 1-β=0.97. For *pezo-1* RNAi (right): Krustal-Wallis one way ANOVA on Ranks, 1-β=0.99. Error bars = SEM, N= 5 animals. EV= **e**mpty RNAi **v**ector (L4440) used as control.

### Swimming is characterized by simultaneous dorso-ventral body wall muscle activation

Swimming behavior was characterized by simultaneous co-activation of dorsal and ventral muscles, with peak activation occurring when the body segment was straight (i.e., at 0.25 and 0.75 cycle fraction) rather than during peak bending (Figure 4A). This pattern differed from the alternating activation observed during crawling, suggesting a distinct neuromuscular strategy for swimming. Also, unlike crawling, where the level of activation was similar between dorsal and ventral muscles, we found that during swimming ventral tail muscles displayed significantly lower activation than tail dorsal muscles. Importantly, reduction of *pezo-1* expression in body wall muscles led to a significant increase in the activation of ventral tail muscles during swimming (Figure 4B). This increase disrupted the typical dorsoventral asymmetry observed in control animals, where ventral tail muscles are normally less active than their dorsal counterparts (Figure 4C).

**Figure 4.**
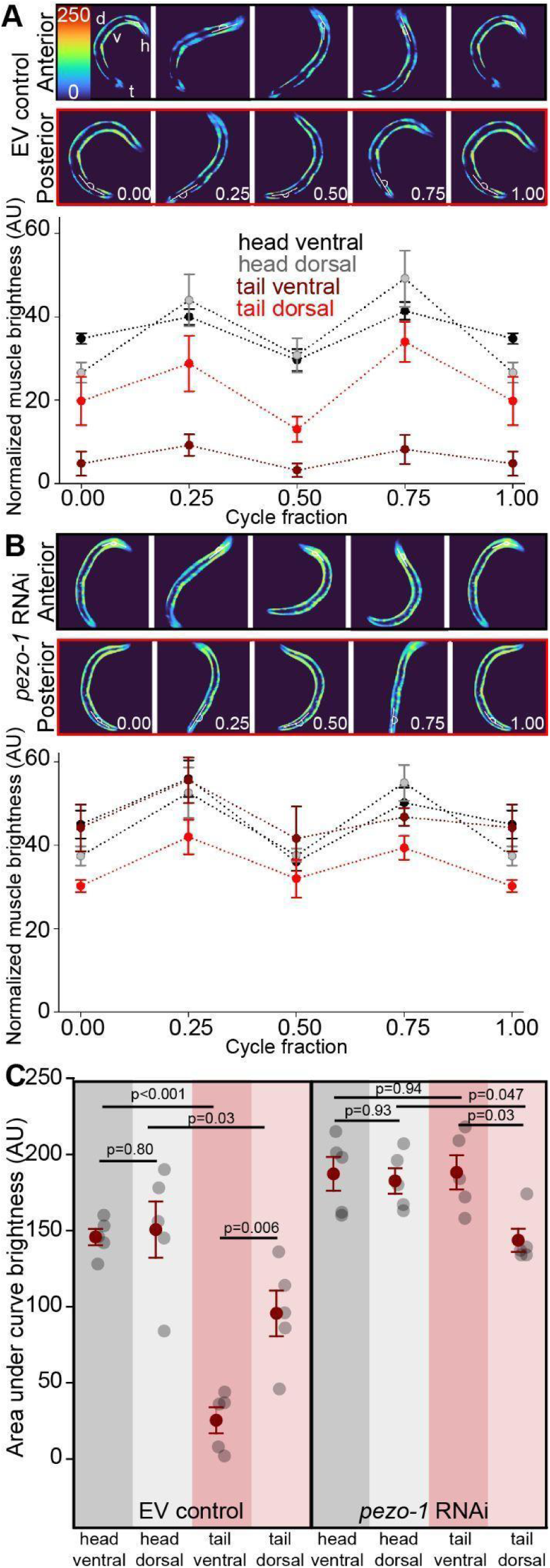
Co-contraction and asymmetrical activation of tail muscles during swimming. **A.** During swimming both dorsal and ventral muscles are activated and relaxed simultaneously. Maximal activation of both ventral and dorsal muscles occurs when the segment is straight. Posterior ventral muscles display reduced activation compared to other muscles. **B.** RNAi-induced reduction in *pezo-1* expression results in relative increase in activation of tail ventral muscles during swimming. **C.** Quantification of area under the curve for A and B shows that tail ventral muscles display significantly lower activation during swimming. RNAi-mediated reduction in *pezo-1* expression causes a significant relative elevation in ventral tail muscle activation. For EV controls (left): One way ANOVA (Holm-Sidak). 1-β=1.0. For *pezo-1* RNAi (right): One way ANOVA (Holm-Sidak). 1-β=0.71. Error bars = SEM, N=5 animals. EV= **e**mpty RNAi **v**ector (L4440) used as control.

### Reduced activation of ventral tail muscles during swimming requires *pezo-1*

To investigate the effects of *pezo-1* knockdown on muscle activation asymmetry, we focused on the differential activation of specific tail muscles. The ventral musculature of *C. elegans* is asymmetrical, with 23 myocytes in its left ventral quadrant and 24 on its right ventral quadrant (Altun and Hall, 2005). We noticed that the myocyte immediately posterior to the spermatheca, ventral myocyte 21 (vm21) appeared to display reduced activation during swimming but participated normally in crawling (Figure 5A, B). We decided to compare vm21 activation to the most posterior myocyte (vm24). In control animals, vm24 was significantly more active than vm21 during swimming, reflecting the lower activation of vm21 (Figure 5D). However, this asymmetry was abolished in *pezo-1* RNAi-treated animals, where vm21 showed increased activation, bringing its activity to levels comparable to vm24 (Figure 5C, E). This was reflected in a significant change in the ratio of vm21/vm24 activation during swimming (Figure 5F). These findings suggest that *pezo-1* is involved in fine-tuning the balance of forces generated by dorsal and ventral tail muscles, which is critical for efficient swimming locomotion.

**Figure 5.**
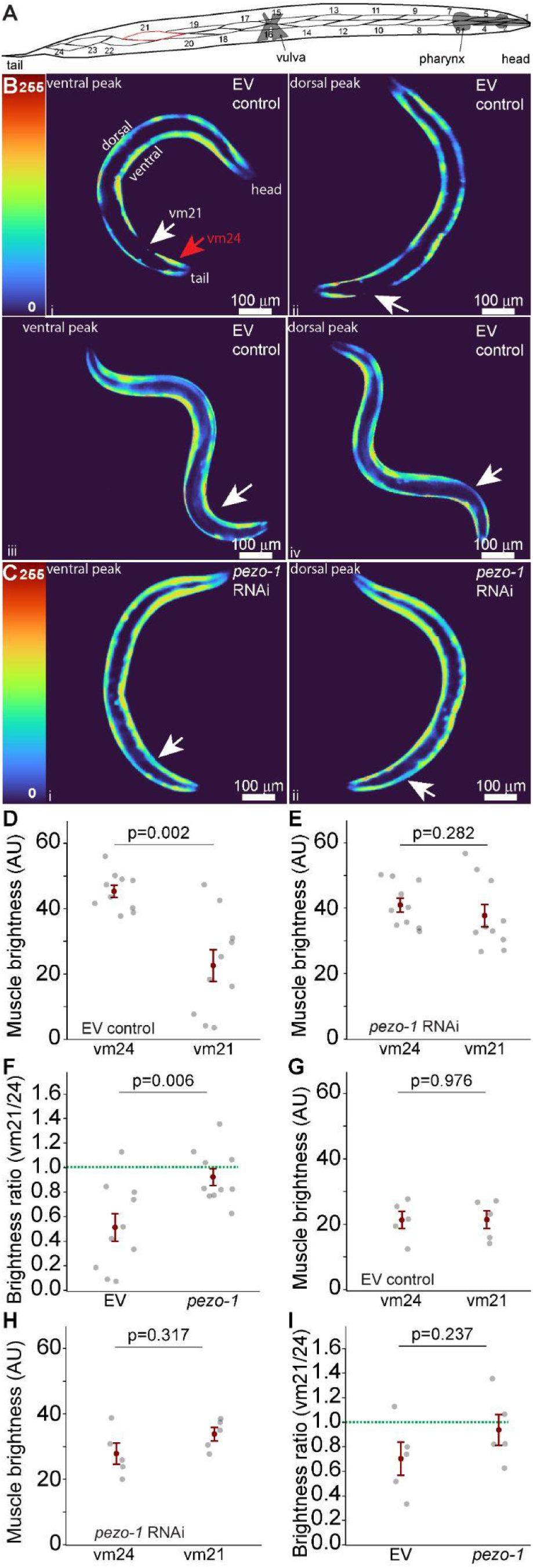
Increased activation of ventral tail muscles during swimming with pezo-1 RNAi. **A.** Diagram, depicting the body wall musculature of the ventral left quadrant of *C. elegans* highlighting vm21 in red. **B.** GCaMP2 activity during swimming showing that ventral myocyte 21 (vm21) is not active at neither the ventral peak (i) nor the dorsal peak of tail contraction during swimming but is active at each of these peaks during crawling (iii and iv respectively. **C.** RNAi-induced reduction in *pezo-1* expression results in activation of vm21 during swimming. White arrows point at vm21, red arrow at vm24. **D.** Average activation of vm21 is significantly lower than vm24 at the peaks of dorsal and ventral contractions during swimming. Paired t-test, 1-β=0.97. **E.** Ventral myocyte 21 and 24 activation during peak ventral and dorsal contraction in *pezo-1* RNAi treated animals during swimming. Paired t-test, 1-β=0.18. **F.** RNAi targeting of *pezo-1* significantly increases the relative activation of vm21 compared to vm24 during swimming. Student t-test, 1-β=0.99. N=10 animals. **G.** Ventral myocyte 21 and 24 activation during peak ventral and dorsal contraction in EV control animals during crawling. Paired t-test, 1-β=0.05. **H.** Ventral myocyte 21 and 24 activation during peak ventral and dorsal contraction in *pezo-1* RNAi treated animals during crawling. Paired t-test, 1-β=0.15. **I.** comparison between vm21/vm24 ratios from EV control and *pezo-1* RNAi treated animals. Student t-test, 1-β=0.91. Error bars = SEM. N=5 animals. EV= **e**mpty RNAi **v**ector (L4440) used as control.

We next performed the same analysis in animals crawling on solid agar surfaces. Unlike swimming, there was no significant difference between the activation of vm21 and vm24 during the peaks on dorsal and ventral contractions (Figure 5G). Similarly to swimming, RNAi targeting of *pezo-1* expression had little effect on vm24 activation but did increase vm21 activation (Figure 5H). However, The vm21/vm24 activation ratio was not different after *pezo-1* RNAi (Figure 5I).

To assess the effects of *pezo-1* downregulation on the dorsal muscles, we compared their activation during swimming and crawling in control and *pezo-1* RNAi treated animals. Downregulating *pezo-1* had no significant effect on dm 21 or dm24 activity during swimming and crawling (Figure S2).

To more closely investigate the role of *pezo-1* in vm21 activation during swimming we quantified GCaMP2 brightness of vm21 in control and *pezo-1* downregulated animals (Figure 6A). We compared the activation of dm21 and vm21 during the ventro-dorsal half of the cycle, and during the dorso-ventral half of the cycle. We found that dm21 was significantly more active than vm21 during each half of the swimming cycle (Figure 6B). However, downregulation of *pezo-1* changed this pattern of activation, with vm21 no longer being significantly less activated than dm21 (Figure 6C).

**Figure 6.**
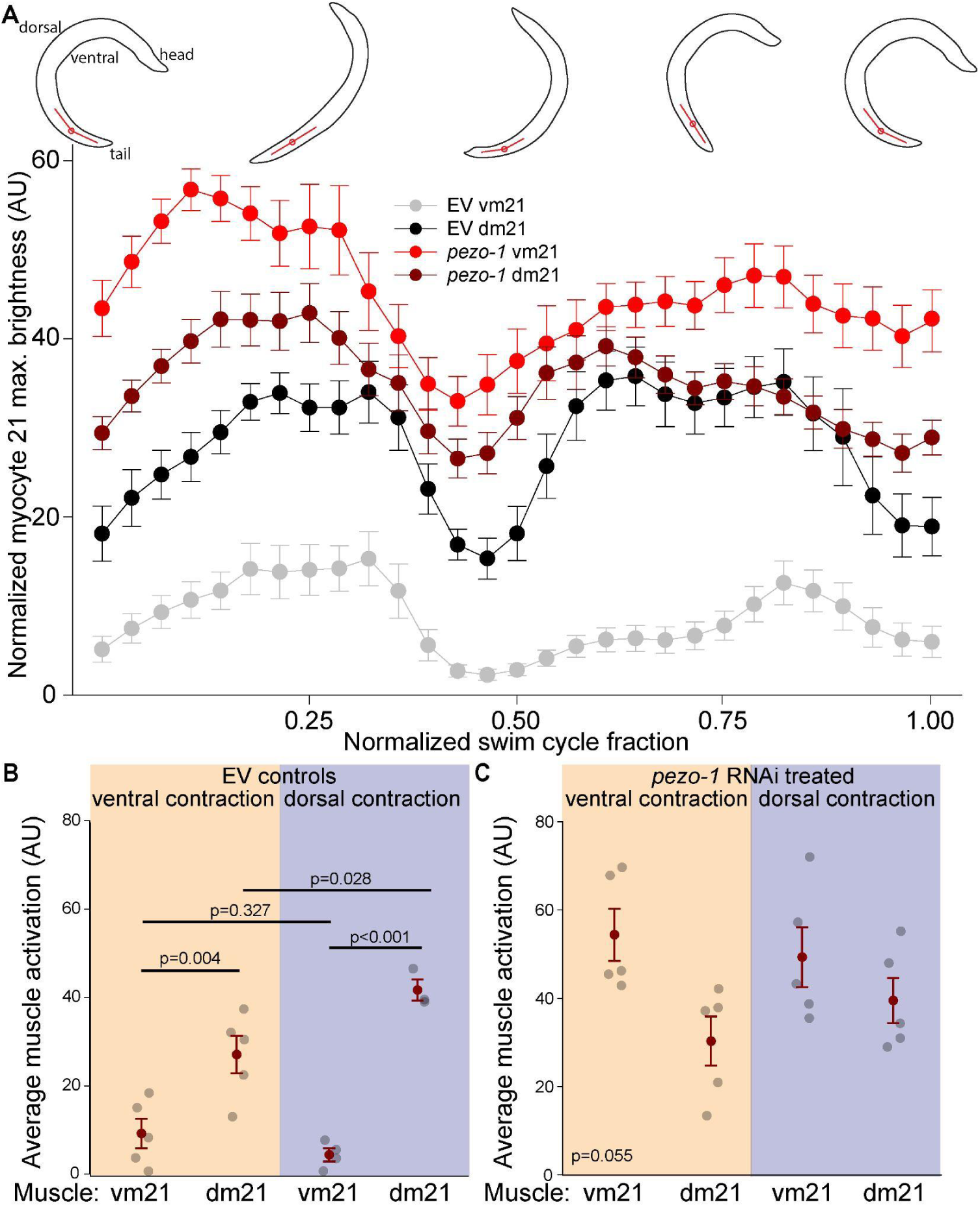
Abolished differential activation between dorsal and ventral tail muscles during swimming with *pezo-1* RNAi. **A.** Activation of ventral and dorsal myocyte 21 (vm21 and dm21) during swimming in control and RNAi treated animals. Red lines indicate the angle of the body segment being measured. **B.** Comparison between ventral and dorsal muscle activation during each half of a swimming cycle shows dm21 are significantly more activated than vm21. One way ANOVA (Holm-Sidak), 1-β=1.0 **C.** *pezo-1* RNAi treatment abolishes the differential activation of dm21 and vm21 during swimming. Kruskal-Wallis One Way ANOVA on Ranks, 1-β=0.75. Error bars = SEM. N=5 animals.

### Dorsoventral asymmetry during swimming requires *pezo-1* expression

Both crawling on solids and swimming in liquids by *C. elegans* are produced by dorsoventral body wall muscle contractions. During crawling, dorsal and ventral contractions are largely symmetrical with similar angular excursions for each. During swimming, dorsal and ventral body contractions are asymmetrical (Pierce-Shimomura et al., 2008). Reflecting this distinction, our results suggest that during swimming muscle activation reflects, and may underlie, the dissimilarity to crawling (Figure 4). Indeed we find that not only is the pattern of muscle activation distinct, but also that the symmetrical contribution of dorsal and ventral muscles seen during crawling is replaced by asymmetrical activation of dorsal and ventral muscles in the tail of the animal (Figure 6). We reasoned that this asymmetrical activation of dorsal and ventral tail muscles would result in the reproduction of asymmetrical tail movement during each half of the swimming cycle.

To test this prediction we measured the curvature of the tail of the worm at m21 during the production of swimming locomotion in control and *pezo-1* RNAi treated animals (Figure 7). In control animals, the rate of change in tail curvature, which is reflected by the slope of the angular excursion, was significantly greater during the ventro-dorsal half of the cycle (labeled “D”) compared to the dorso-ventral half (labeled “V”) (Figure 7A, B). However, downregulation of *pezo-1* significantly reduced the rate of curvature change during the ventro-dorsal phase, making it comparable to the dorso-ventral phase (Figure 7B). This decrease in the rate of curvature change was accompanied by a significant decrease in the angular excursion of the tail for *pezo-1* downregulated animals (Figure 7C).

**Figure 7.**
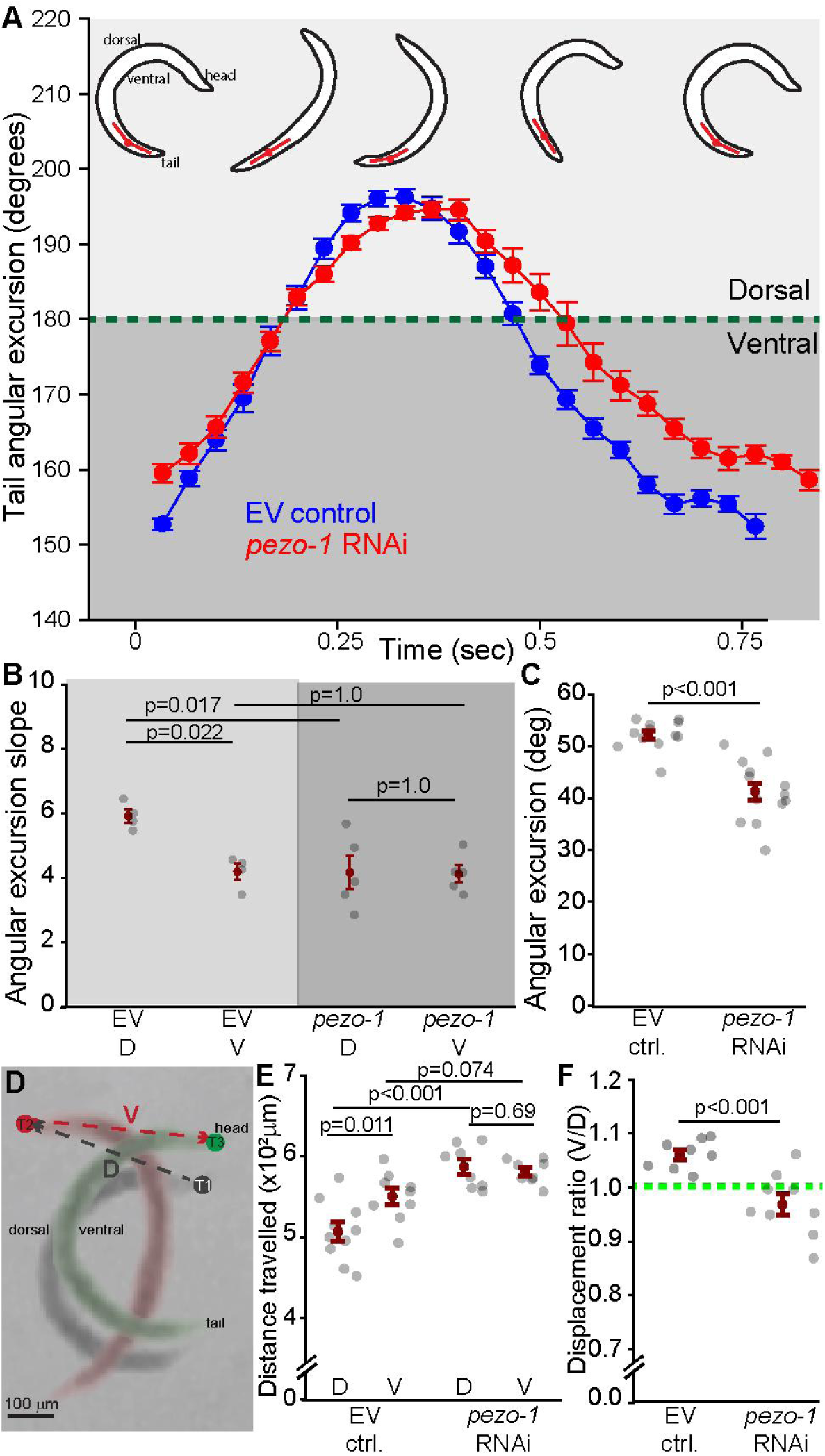
Decreased angular excursion of the tail during swimming with *pezo-1* RNAi. **A.** Average angular excursion of the tail during swimming (blue = control, red = *pezo-1* RNAi). Most of the tail range of motion during swimming is ventral (below the green dotted line). N=4 animals. **B.** Quantification and comparison of the value of the angular excursion slopes during ventro-dorsal contraction (D), and dorso-ventral contraction (V) in control (L4440) and *pezo-1* RNAi silenced animals. In control animals the angular excursion during the first (D) half of the cycle has a significantly greater slope than during the second (V) half. This asymmetry is abolished in RNAi-treated animals where a decrease in the ventro-dorsal slope causes both halves to display symmetric movements. One way ANOVA (Holm-Sidak). 1-β=0.79. **C.** RNAi-induced reduction in *pezo-1* expression results in a significant decrease in the angular excursion of the tail during swimming. Student t-test. 1-β=1.0. N=14 animals **D.** Composite image showing the ventro-dorsal (D) and dorso-ventral (V) displacement of the head of a worm during the production of one swimming cycle. **E.** Control animals displayed a grated displacement during the V half of the cycle, but this difference was abolished in *pezo-1* RNAi treated animals. One way ANOVA (Holm-Sidak). 1-β=1.0. N=10 animals **F.** Comparison of lateral displacement ratio between control (EV) and *pezo-1* RNAi treated animals. Welch’s t-test, 1-β=1.0. Error bars = SEM. N=10 animals. EV= **e**mpty RNAi **v**ector (L4440) used as control.

To determine if these curvature changes affected the performance of the locomotor behavior we quantified the lateral displacement of the head of the worm during swimming for both control and *pezo-1* RNAi treated animals (Figure 7D). We found that in control animals lateral displacement was significantly greater during the dorso-ventral half of the cycle (Figure 7E, consistent with an effective propulsion of the worm). However, *pezo-1* RNAi treatment increased lateral displacement during the ventro-dorsal half of the cycle abolishing the difference between the two halves of the cycle (Figure 7F).

### Downregulation of *pezo-1* leads to altered motor output

Kinematic analysis of swimming behavior using Tierpsy Tracker revealed that downregulation of *pezo-1* significantly altered the most commonly performed postures, as represented by eigen projections (Figure S3). Specifically, *pezo-1* knockdown resulted in significant changes in the contributions of eigen projections 3, 5, and 6, indicating alterations in the worm’s swimming posture (Figure S3B-D). These changes likely reflect the disrupted muscle activation patterns and reduced tail curvature observed in *pezo-1* RNAi-treated animals. Path curvature, a measure of the worm’s swimming trajectory, was significantly reduced in *pezo-1* knockdown animals, further indicating impaired swimming performance (Figure 8).

**Figure 8.**
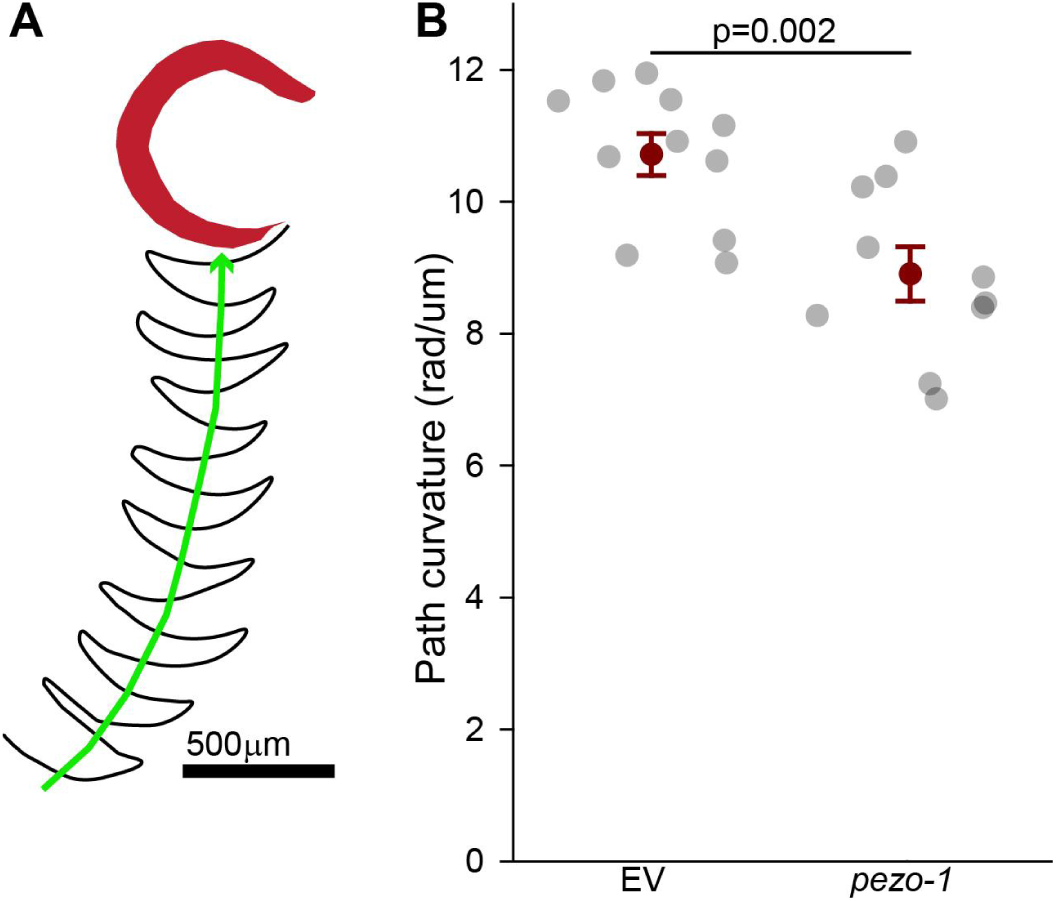
Reduction in path curvature during swimming *pezo-1* RNAi. **A.** A schematic representation of the path of tail tip of swimming *C. elegans*. **B.** Quantification of path curvature (rad/um) comparing control (EV) and *pezo-1* RNAi silenced animals. Path curvature is defined as the angle of the worm’s path divided by the distance traveled. *pezo-1* RNAi silenced animals exhibited significantly reduced path curvature, meaning less curved movement. Student t-tests. 1-β=1.0. Error bars = SEM. N>10 animals per each (datapoint) assay. EV= **e**mpty RNAi **v**ector (L4440) used as control.

### Tail Muscles Exhibit Differential Expression of Dystrophin

The dramatic effects of *pezo-1* knockdown on the swimming behavior (Figure 2, 8, S3) are consistent with the increased activation of the vm21 muscle in pezo-1 RNAi treated animals (Figure 5). Taken together, this suggests that vm21 plays a crucial role in this behavior, and, because PEZO-1 is a mechanoreceptor, that vm21 may be under particular mechanical strain during swimming. This indicates that vm21 membrane stability is crucial and key structural proteins should be expressed in abundance. We tested this idea by examining the expression of dystrophin, a key structural protein, in these muscles using a transcriptional reporter strain. We found that vm21 exhibited significantly higher expression of dystrophin compared to neighboring muscles such as vm20 (Figure 9A and B). This differential expression suggests that vm21 may have a specialized role in tail movement during swimming, potentially contributing to the muscle’s structural integrity and responsiveness to mechanical stress.

**Figure 9.**
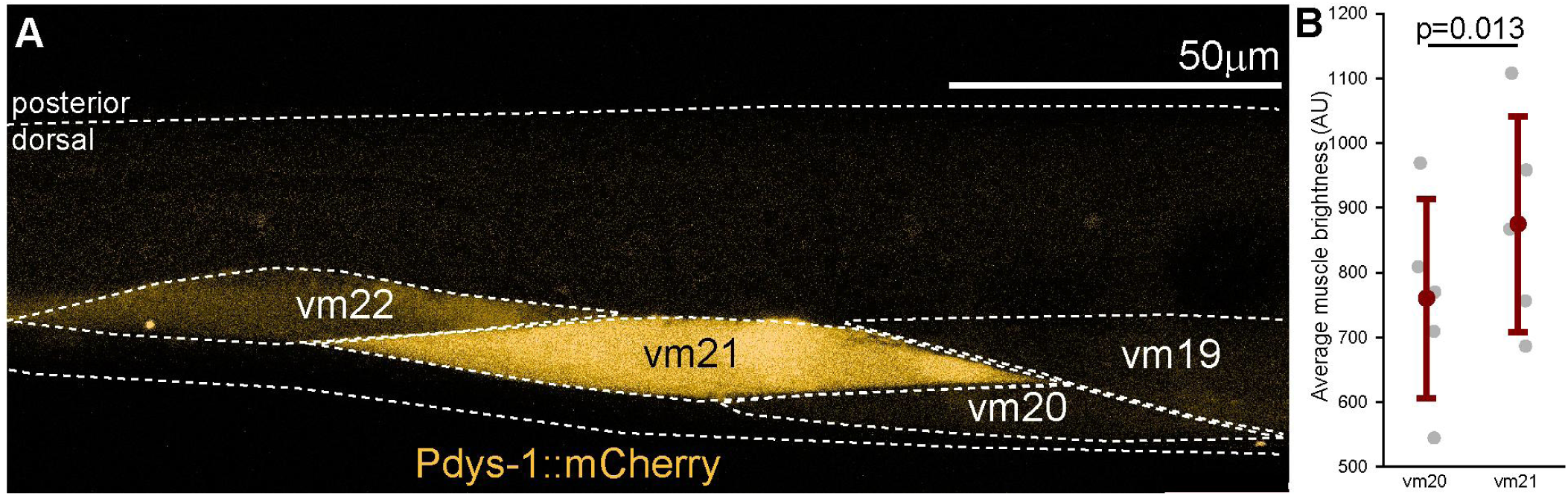
Increased dystrophin expression in ventral myocyte 21 (vm21). **A.** Transcriptional reporter (mCherry) for dystrophin (*dys-1*) shows increased expression in vm21 compared to nearby muscles like vm20. **B.** Average mCherry brightness of vm21 is significantly greater than vm20. Paired t-test. 1-β=0.87. Error bars = Standard deviation. N=5 animals.

Please refer to files Supplementary file 1 for a spreadsheet containing the summary and comparative statistics for the data presented in this study and to Supplementary file 2 for a spreadsheet containing the raw measurements used in the study.

## DISCUSSION

Our study investigated the role of the mechanoreceptor PEZO-1 in the body wall musculature of *C. elegans* using a combination of high-throughput kinematics, genetically encoded calcium indicators, and RNAi. The results demonstrate that PEZO-1 has distinct roles in modulating muscle activity in swimming and crawling.

### Muscle Activation and Locomotion

As anticipated based on prior research, we observed differences in muscle activation patterns between crawling and swimming. Crawling relies on the symmetrical, alternating activation of antagonistic dorsal and ventral muscles, with peak activation coinciding with the peak of body bending (Figure 3; Pierce-Shimomura et al., 2008). In contrast, swimming exhibited a fundamentally different pattern. We found that antagonistic muscles were continuously co-activated, with peak activation occurring when the body segment was straight and minimal activation when the segment was maximally bent (Figure 4). This continuous co-activation and its asymmetrical nature suggest a unique adaptation for movement in a liquid environment, as opposed to the solid substrate of crawling.

These findings are consistent with the concept of the NMT, where the relationship between motor neuron firing patterns and muscle contraction is nonlinear and dependent on the context of the behavior (Brezina et al., 2000a, b; Brezina and Weiss, 2000). The shift in activation patterns between crawling and swimming further supports the idea that different locomotor strategies require distinct neuromuscular control mechanisms (Vidal-Gadea et al., 2011).

### Differential Role of *pezo-1* in Locomotion

Our data revealed that PEZO-1 is differentially involved in swimming versus crawling. The reduction in *pezo-1* expression resulted in altered body postures during swimming, including significant changes in eigen projections 3, 5, and 6 (Figure S3), but had a minimal effect during crawling (Komandur et al., 2023). Specifically, downregulating *pezo-1* significantly impacted the swimming path curvature and the eigen projections that capture the most commonly performed postures (Figures 8 and 9). This suggests that PEZO-1 is particularly important for fine-tuning muscle activity in response to the mechanical demands of swimming.

The observed differential function of PEZO-1 aligns with previous studies highlighting the importance of mechanoreceptors in adaptive locomotion (Gjorgjieva et al., 2014). Mechanoreceptors like PEZO-1 may provide a mechanism for muscles to adjust their contractile properties in response to changes in external forces, thereby contributing to the flexibility and adaptability of movement.

### Mechanisms of *pezo-1* Modulation of Muscle Activation

Our finding that RNAi-mediated reduction of *pezo-1* expression leads to increased muscle activation, particularly the asymmetric increase in ventral tail muscle activation during swimming (Figures 5 and 6), is counterintuitive given that PEZO-1 is a mechanosensitive cation channel likely involved in calcium influx. This raises the question of how a reduction in PEZO-1 could lead to increased intracellular calcium levels and heightened muscle activity.

Several mechanisms could explain our observations. One mechanism suggests that PEZO-1 channels may be asymmetrically distributed across dorsal and ventral muscles, with a higher concentration in dorsal muscles. This asymmetry might contribute to greater calcium influx and activation in dorsal muscles during swimming but not crawling conditions. When *pezo-1* expression is reduced, this differential influx is lost, potentially triggering a compensatory increase in calcium release from intracellular stores, such as the sarcoplasmic reticulum. This could particularly affect ventral muscles. This could lead to the observed increase in ventral muscle activation and loss of dorsoventral asymmetry during swimming. The differential expression of *pezo-1* in posterior muscles, as suggested by transcriptomic data (Ghaddar et al., 2023), supports this model.

Alternatively, reduced activation of dorsal tail muscles brought about by a reduction of *pezo-1* expression in those muscles could trigger a compensatory global increase in neuromuscular activation. This global increase would explain both the loss of asymmetrical activation between dorsal and ventral tail muscles and the observed increase in intracellular calcium levels across all body wall musculature. The idea that a reduction in local muscle activation after *pezo-1* knockdown could result in a systemic response is consistent with known feedback mechanisms where decreased force generation in one muscle group leads to a compensatory increase in overall neuromuscular activity to maintain locomotor function (McLean and Goudi, 2004).

Another potential explanation involves neural pathways. PEZO-1 might influence the neural circuits controlling muscle activation. While body wall muscles are readily susceptible to RNAi by feeding, most neurons in *C. elegans* are refractory to this treatment, including those innervating body wall muscles (Calixto et al., 2010; Firnhaber and Hammarlund, 2013). *Pezo-1* is expressed in GABAergic neurons, which inhibit muscle contraction. A loss of PEZO-1 in these neurons could decrease inhibitory signaling, leading to increased muscle activity, particularly in the ventral muscles during swimming. However, the specificity of RNAi by feeding to muscle tissue, confirmed by the literature (Calixto et al., 2010; Firnhaber and Hammarlund, 2013), suggests that this mechanism might be less likely unless there is an indirect effect mediated by muscle-nerve communication.

### Implications for Neuromuscular Control and Behavioral Plasticity

The differential activation of tail muscles during swimming, with a significant increase in the activation of ventral tail muscles in *pezo-1* RNAi-treated animals (Figure 6), suggests that PEZO-1 may play a role in modulating the balance of forces generated by dorsal and ventral muscles. This balance is crucial for maintaining the efficiency and directionality of swimming, as the loss of this asymmetry led to slower swimming speeds in PEZO-1-deficient worms (Figure 2).

These findings highlight the potential of PEZO-1 and similar mechanoreceptors to act as modulators of neuromuscular control, enhancing the behavioral repertoire of organisms without necessitating more complex neural circuitry. This concept is particularly relevant in the context of evolutionary biology, where the ability to adapt movement patterns to different environments can provide a selective advantage (Pierce-Shimomura et al., 2008; Vidal-Gadea et al., 2011).

### Limitations and Future Directions

While our study provides insights into the role of PEZO-1 in *C. elegans* locomotion, it is not without limitations. One limitation is the use of RNAi by feeding, which may not achieve complete knockdown of gene expression, particularly in certain tissues. This partial knockdown might result in residual PEZO-1 activity that could influence the observed outcomes. Additionally, our experiments relied on the differential RNAi refractory nature of muscles and neurons to target the muscles rather than the neurons that innervate them. It is possible that some neurons displaying higher than normal susceptibility to RNAi by feeding might have been affected by our treatment. However, previous work established that none of the motor neurons innervating body wall muscles (which include excitatory cholinergic and inhibitory GABAergic neurons) are targetable by RNAi by feeding unless they are made to express an ectopic dsRNA transporter channel (Firnhaber and Hammarlund, 2013).

Future research could address these limitations by using tissue-specific or inducible RNAi systems to achieve gene knockdown in specific myocytes or neurons. Furthermore, rescue experiments that reintroduce PEZO-1 selectively in either muscle or neural tissue would help clarify the relative contributions of these tissues to the observed phenotypes. Another area for future investigation is the potential interaction between PEZO-1 and other mechanosensitive channels or structural proteins, such as dystrophin, to better understand the broader mechanotransduction network in muscle function.

In conclusion, our findings underscore the importance of PEZO-1 in modulating muscle activity during swimming in *C. elegans*, demonstrating that mechanoreceptor expression in muscles is a critical factor in adaptive locomotion. This study contributes to the broader understanding of neuromuscular control and highlights the potential for mechanoreceptors to enhance behavioral plasticity in response to environmental demands.

## METHODS

### Strains and Cultivation

*Caenorhabditis elegans* strains were maintained on nematode growth medium (NGM) agar plates seeded with Escherichia coli OP50 as a food source, following standard protocols (Brenner, 1974). The strains used in this study included N2 (wild type), ZW495 (*myo-3p::GCamp2*), COP1367 (*pezo-1*(IV:9366441)), and AG405 (*pezo-1* C-terminus deletion of Exon 27-33 + introns). Strain COP1367 was a kind gift from Dr. Valeria Vásquez, and strain AG405 was provided by Dr. Andy Golden and Dr. Xiaofei Bai. Unless otherwise noted, animals were cultivated at 20–22°C with 30–37% humidity.

### Transgenic Reporter Construction

To construct transcriptional reporter strains, we PCR-amplified approximately 4kb upstream promoter regions of the *dys-1* gene from N2 genomic DNA. The amplified products were inserted into a transcriptional reporter vector containing mCherry downstream of the promoter, followed by the unc-54 3’ UTR sequence. Transgenic strains were generated by microinjecting these constructs into the gonads of young adult hermaphrodites. Positive transgenic animals expressing mCherry were identified by fluorescence microscopy, and stable lines were established through propagation.

Expression patterns were analyzed using confocal microscopy, and the quantification of *dys-1* expression was done using ImageJ.

### Gene Knockdown by RNAi

RNA interference (RNAi) was performed using the feeding method described by Conte et al. (2015). Briefly, day 1 adult worms were allowed to lay eggs on RNAi plates seeded with IPTG-induced HT115 bacteria carrying either the L4440 empty vector (control) or *pezo-1* dsRNA construct. Eggs were incubated at 20°C until they developed into L4 larvae. RNAi by feeding is effective in non-refractory tissues, particularly body wall muscles, and targets genes expressed within these tissues (Grishok, 2005). For controls, animals were treated with bacteria carrying the empty L4440 vector.

### Tissue Fixation and Staining

Worms were collected from NGM plates using liquid NGM and washed three times to remove excess bacteria. Fixation was performed in PBS containing 16% formaldehyde for 15 minutes at room temperature, followed by three washes with 1X PBS. The freeze-cracking method described by Duerr (2013) was employed to prepare samples for staining. For dual staining, samples were incubated overnight at 4°C with a mixture of Phalloidin conjugated to iFluor 488 and Wheat Germ Agglutinin (WGA) conjugated to CF680 to visualize actin filaments and cellular membranes. Following staining, samples were washed three times with 1X PBS before imaging.

### Confocal Imaging

Confocal microscopy was performed using a Leica SP8 confocal microscope equipped with Lightning mode. Z-stacks were acquired at regular intervals through the specimen depth using a HC PL APO CS2 63x/1.40 oil immersion objective. Images were acquired using sequential imaging to avoid crosstalk between channels. Leica Application Suite X software was used for image acquisition and post-processing, including standard brightness and contrast adjustments to optimize image visualization while maintaining data integrity.

### Behavior Analysis

The swimming and crawling behaviors of *C. elegans* were analyzed using the Tierpsy Tracker software (Javer et al., 2018). Videos of individual worms were recorded at 10 frames per second for approximately 17 seconds. More than 10 worms were analyzed per condition. Tierpsy Tracker was used to quantify crawling and swimming speeds, path curvature, and body postures. Kinematic analysis was performed to identify the most common postures (eigen projections), and path curvature was calculated based on the centroid’s side-to-side displacement relative to forward progress.

### Calcium Imaging

Calcium dynamics in body wall muscles were measured using GCaMP2 expressed under the control of the *myo-3* promoter. Worms were illuminated using an X-cite 120 PC Q light source, and calcium transients were recorded. Fluorescence intensity was measured using ImageJ, with regions of interest defined around specific muscles. To assess relative calcium levels, the fluorescence intensity of GCaMP2 was normalized against background fluorescence measured in adjacent areas devoid of muscle tissue.

Additionally, for tail muscle brightness analysis during swimming behavior, fluorescence intensity data were normalized to the duration of each swim cycle to account for variability in cycle length. The normalization was done by first identifying the longest swim cycle duration among all recorded cycles. The duration of each cycle was then scaled relative to this maximum duration. Fluorescence intensity values were interpolated to match the normalized time points of each cycle.

To quantify average muscle activation, the GCaMP2 brightness data of the tail muscle during each swim cycle were analyzed by dividing the cycle into two parts (the first half and the second half). The average GCaMP2 brightness was calculated separately for each part of each cycle. Then, these averages were averaged across all cycles to obtain a mean value for each parts per animal. This final average provides an overall measure of muscle activation for each animal during the observed swim cycles.

### Tail Angle and Lateral Displacement Measurements

Tail angle changes during swimming were analyzed using the same recordings used for calcium imaging. The angle at the tail gap was calculated by measuring the angle between the tail tip and a fixed anterior point along the body midline for each frame. Angular excursion and curvature dynamics were plotted over time, and slopes were analyzed to determine the rate of angular change.

Lateral displacement was measured by analyzing the movement of the worm’s head during swimming. Displacement was quantified by measuring the distance between the head’s position during the maximum ventral bend and the maximum dorsal bend in the swimming cycle. ImageJ was used to calculate the displacement distance, which provided insights into the worm’s locomotor strategies.

### Statistical analysis

Statistical analyses were conducted using SigmaPlot 14.5. Data were assessed for normality using the Shapiro-Wilk test and for equal variance using the Brown-Forsythe test. Outliers were identified using the interquartile range method and removed from analysis. Paired parametric datasets were compared using paired t-tests, while unpaired datasets were compared using Student’s t-tests if groups had equal variance, or Welch t-tests if groups failed the equal variance test. For comparisons involving more than two datasets, One-Way ANOVA was used, followed by Holm-Sidak post hoc tests. Non-parametric data were analyzed using the Rank Sum test or ANOVA on Ranks followed by Dunn’s post hoc tests. Statistical significance was set at an alpha level of 0.05 and a beta of 0.20. Statistical power (1-β) was calculated in SigmaPlot for parametric data. For non-parametric tests power was calculated using the Monte Carlo Simulation method. Detailed descriptive and comparative statistics for each figure along with raw data are available in the supplementary files. Video files are accessible through Git-Hub (Fazyl and Vidal-Gadea, 2024).

## ACKNOWLEDGEMENTS

The COP1367 strain was a kind gift from Dr. Valeria Vásquez. The AG405 strain was a kind gift from Dr. Andy Golden and Dr. Xiaofei Bai. Other strains were provided by the Caenorhabditis Genetics Center (funded by NIH Grant P40 OD010440). Funding was provided by the National Insitutes of Health, National Institute of Arthritis and Musculoskeletal and Skin Diseases award 2R15AR068583-02, and the National Science Foundation, Division of Molecular and Cellular Biosciences award 1818140 to A.G.V.-G.

## Figures

**Figure S1.**
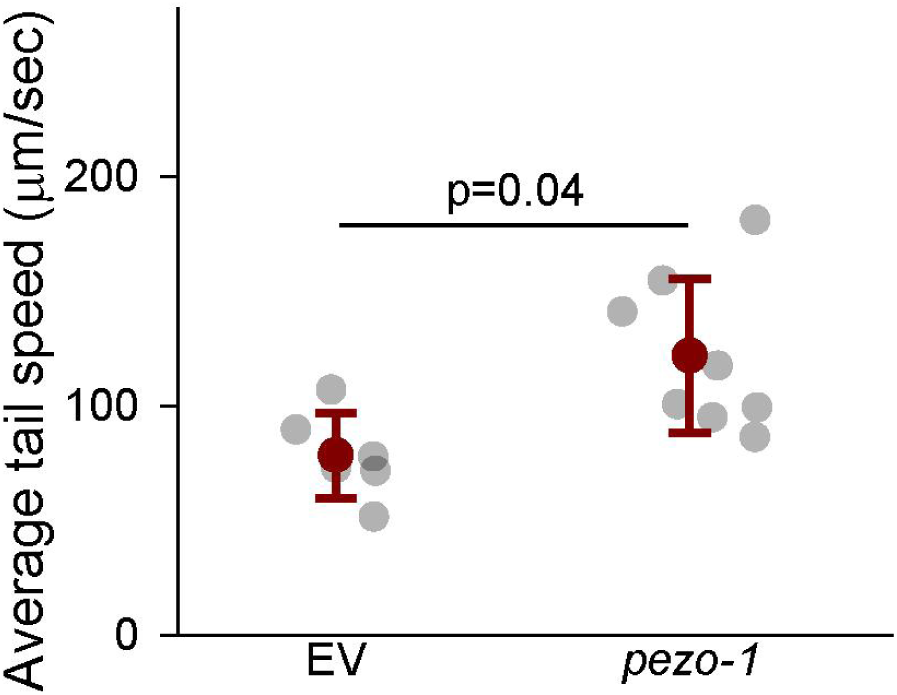
Increased crawling speed with *pezo-1* RNAi. RNAi targeting *pezo-1* expression increases the average speed of the tail during crawling compared to EV= **e**mpty RNAi **v**ector (L4440) used as control. Student t test. 1-β=0.49, Error bars = SEM. N>10 animals per each (datapoint) assay.

**Figure S2.**
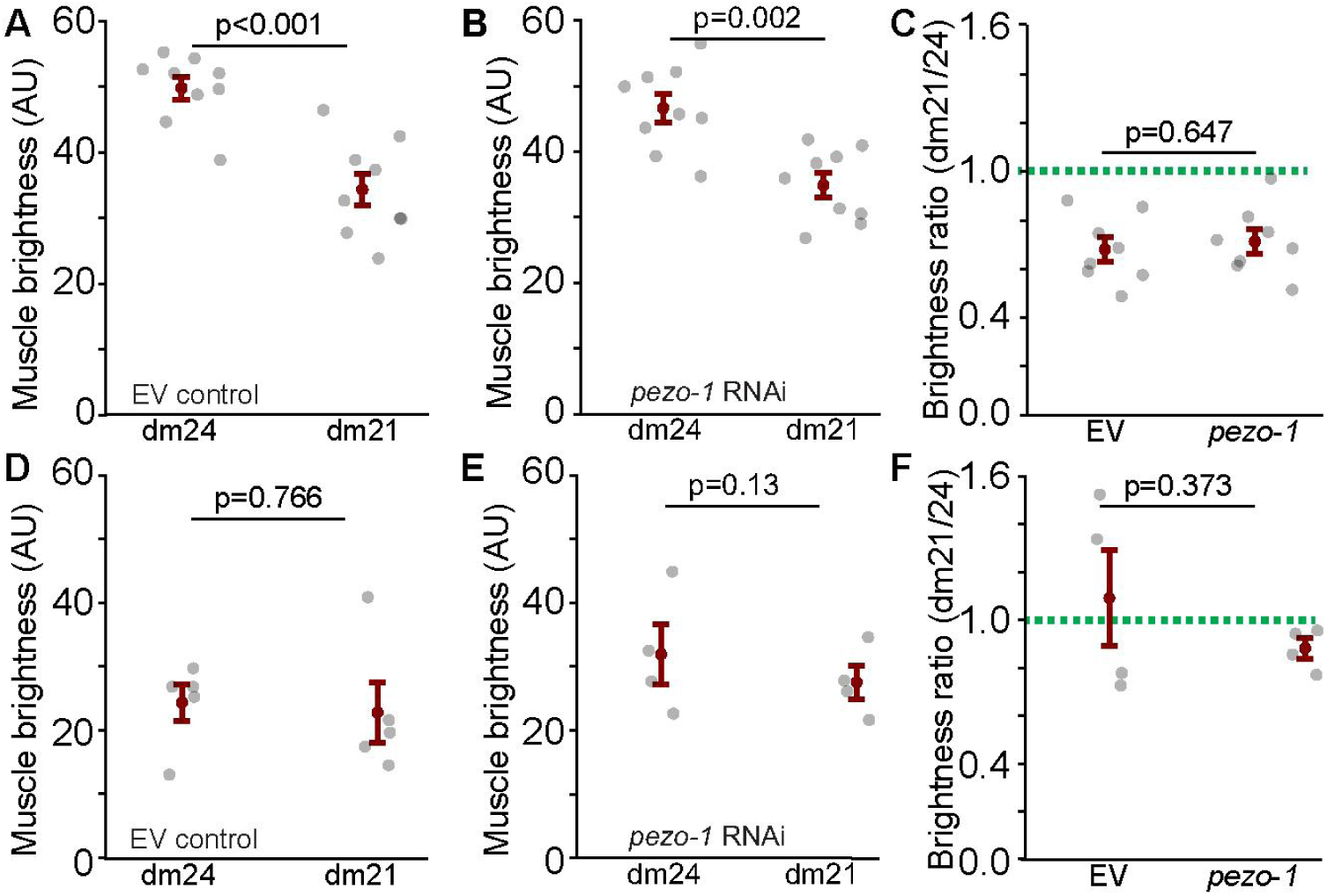
No selective increase in dorsal tail muscle activation during swimming with *pezo-1* RNAi. **(A)** Dorsal myocyte 21 and 24 activation during peak ventral and dorsal contraction in EV control animals during swimming. Paired t-test. 1-β=1.0. **(B)** Dorsal myocyte 21 and 24 activation during peak ventral and dorsal contraction in *pezo-1* RNAi treated animals during swimming. Paired t-test. 1-β=0.97. **(C)** Quantification shows no significant effect on relative activation of dm21 vs. dm24 with *pezo-1* RNAi during swimming peak tail bending. Welch’s t-test. 1-β=0.83. N=10 animals. **(D)** Dorsal myocyte 21 and 24 activation during peak ventral and dorsal contraction in EV control animals during crawling. Paired t-test. 1-β=0.06. **(E)** Dorsal myocyte 21 and 24 activation during peak ventral and dorsal contraction in *pezo-1* RNAi treated animals during crawling. Paired t-test. 1-β=0.3. **(F)**. Quantification shows no significant effect on relative activation of dm21 vs. dm24 with *pezo-1* RNAi during crawling peak tail bending. Welch’s t-test. 1-β=0.89. N=5 animals. EV= **e**mpty RNAi **v**ector (L4440) used as control. Error bars = SEM.

**Figure S3.**
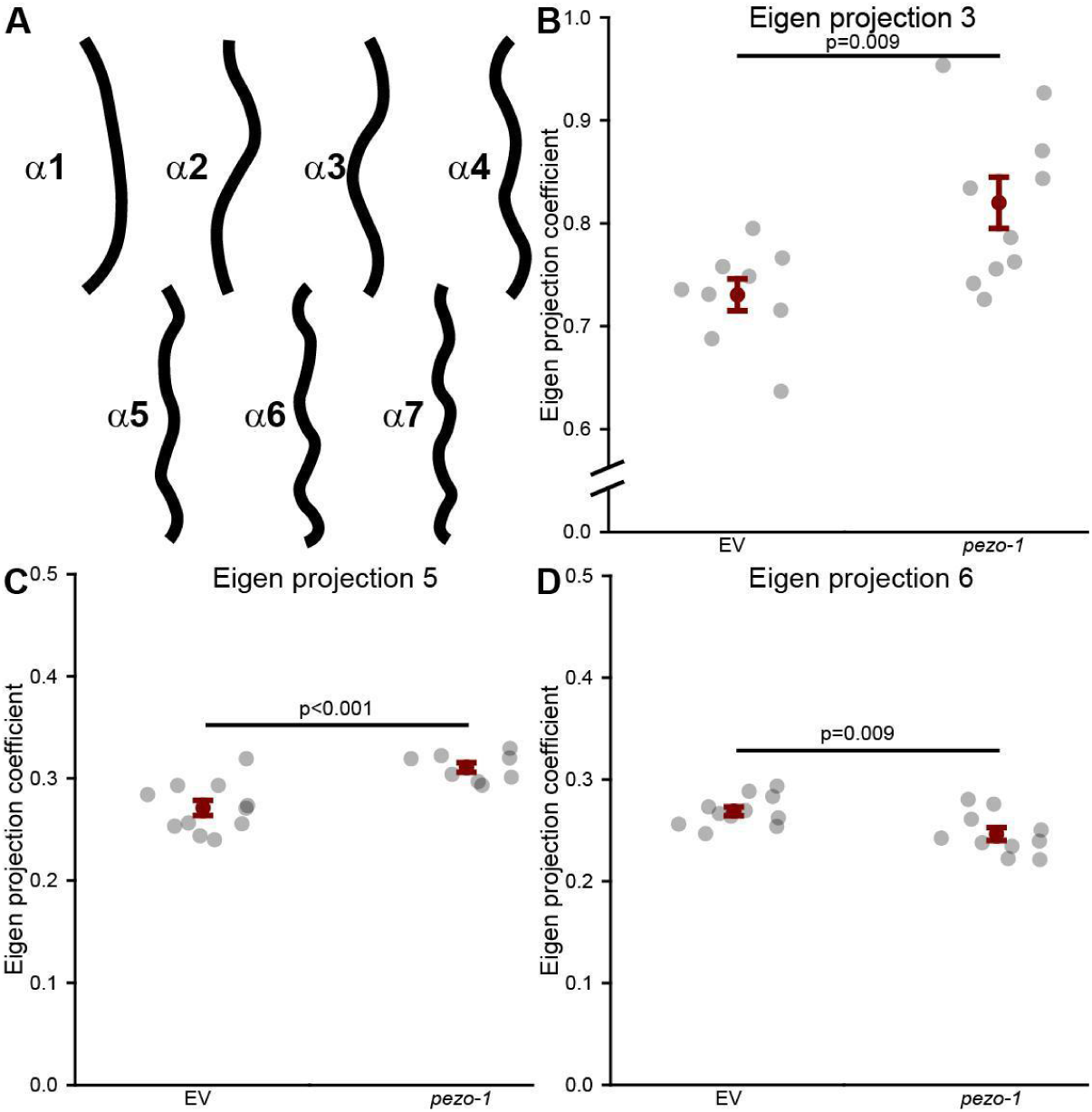
Alteration in swimming posture with *pezo-1* RNAi. **A.** Kinematic analysis performed using Tierpsy extracts the most common postural components of the animal behavior during swimming (eigen projections). RNAi targeting of *pezo-1* expression significantly impacted the eigen coefficients or projections 3, 5, and 6, resulting in a significant increase in contributions of projections 3 (**B**) and 5 (**C**), and a significant decrease in projection 6 (**D**). EV= **e**mpty RNAi **v**ector (L4440) used as control. Student’s t-tests. 1-β>0.99. Error bars = SEM.

